# Detection of latent Epstein-Barr virus gene expression in single-cell sequencing of peripheral blood mononuclear cells

**DOI:** 10.1101/2022.05.24.492331

**Authors:** Alan Bäckerholm, Yarong Tian, Isak Holmqvist, Guojiang Xie, Diana Vracar, Sanna Abrahamsson, Carolina Guibentif, Ka-Wei Tang

## Abstract

Epstein-Barr virus (EBV) DNA is regularly found in the blood of patients with EBV associated diseases and occasionally in healthy individuals. However, EBV infected primary B-lymphocytes have not yet been detected using scRNA seq. Here, we screened the viral transcriptome in single cell RNA sequencing datasets from peripheral blood to identify virus infected cells. Whereas EBV RNA was detected in an immunocompromised patient, EBV associated nasopharyngeal carcinoma and multiple sclerosis samples did not display any levels of circulating EBV RNA. We further screened whole-blood samples from a cohort of immunosuppressed patients for viral transcripts using a custom enhanced RT-qPCR panel and detected latency programs dominated by noncoding RNAs (EBERs and RPMS1). To explore the interplay between the EBV and the host-cell transcriptome profile, we used enriched B-lymphocytes from a splenectomy patient with 30% EBER positivity estimated by *in situ* hybridization and performed 5’ single-cell RNA sequencing with paired VDJ profiling. The EBV expression pattern of the patients’ B-lymphocytes confirmed the RT-qPCR assay with RPMS1 and LMP-1/BNLF2a/b significantly dominating the sequenced EBV polyadenylated RNA. A comparison between the expression profile of EBV positive B-lymphocytes and healthy controls B-lymphocytes revealed the upregulation in genes involved in cell population proliferation when infected with EBV. This is further supported by a measurable polyclonal expansion in the patient, as compared to a control, emphasizing EBV’s role in a host-cell’s tendency for cellular expansion. However, when contrasting to cells that have undergone malignant transformation, the primary EBV infected cells display a rather dissimilar expression profile, even to cells that are supposed to simulate primary EBV infection (I.e. Lymphoblastoid Cell Lines). This implies that during primary infection of EBV, the host-cell enters a state of premalignancy rather than a complete oncogenic transformation at the initial time of infection

## Introduction

The capability of viruses to remain in a dormant state within particular host cells is a hallmark feature of members of the *Herpesviridae*. Whereas the α-herpesviruses persist in neuronal cells, the latent reservoirs of β- and γ-herpesviruses are predominantly situated in leukocytes and can be readily detected in peripheral blood. In whole-genome sequencing data of healthy blood samples, Epstein-Barr virus (EBV) and cytomegalovirus DNA, were detected in 14.47% and 0.35% of samples, respectively^1^. Approximately, 1 in 100,000-1,000,000 circulating B-lymphocytes contain EBV^2^. The sparse viral gene expression during latency is imperative for immune evasion and poses therapeutic challenges. This is particularly relevant with respect to EBV and its inherent transforming capacity coupled to latent infection. Besides the self-limiting lymphoproliferation seen in infectious mononucleosis, EBV is associated with various lymphoid and epithelial malignancies.

The inability to distinguish the cell type in which the infectious particle resides is a material limitation of the bulk-sequencing approach. This issue can, however, be addressed by utilizing single-cell RNA sequencing (scRNA-seq). It has been shown that cytomegalovirus transcripts can be detected using scRNA-seq^3^. However, the limited number of identified cells hampers transcriptional profiling of infected cells. Furthermore, identification of viral reads in the main cell type reservoir for latent herpesvirus infections has not been accomplished using scRNA-seq. The characterization of the infected cell is pertinent to understand the pathogenic mechanisms and to determine the tropism of persistent viral etiologies.

B-lymphocytes are the main reservoir of latent EBV and the majority of associated lymphoid neoplasms often arise from this cell type, however rare lymphomas may originate from NK-cells or T-lymphocytes^4^. EBV-infected cells and neoplasms are believed to express a limited set of EBV-encoded proteins and noncoding RNAs in specific latency programs. The EBV nuclear antigens (EBNAs) and latent membrane proteins (LMPs), in concert with the noncoding EBV small RNAs (EBERs) and *Bam*-HI A rightward transcripts (BART) microRNAs, are believed to constitute the core that maintains latency^5^. EBV gene expression has been explored at the single-cell level in the epithelial cells of nasopharyngeal carcinoma^6,7,8, 9^. Similarly, the transcriptome of EBV-transformed lymphoblastoid cell lines has also recently been investigated using scRNA-seq^10^. However, viral reads have not been previously reported from primary hematological tissue by scRNA-seq. Unlike neoplastic cells, primary EBV-infected lymphocytes likely retain a constitutional genome and the perturbed transcriptome would thus reflect adaptations solely as a response from the viral elements.

In this study, we have optimized virus detection in peripheral blood mononuclear cells (PBMC) using scRNA-seq. An initial large screen of publicly available scRNA-seq datasets from PBMCs confirmed that virus detection is possible, albeit with low sensitivity. In contrast, by identifying specific patient cohorts we consistently detected latent EBV gene expression and identified samples suitable for scRNA-seq. With the addition of cell type enrichment an abundant number of primary EBV-infected B-lymphocytes were detected using scRNA-seq. By adapting standard bioinformatics pipelines to accommodate for the EBV genome and complementing with VDJ linkage sequences we were able to increase the classification of EBV-positive cells almost eight-fold and identify clonotypes with specific cellular markers for primary latently EBV-infected B-lymphocytes. Finally, we confirm and extend EBV’s role in cellular expansion^11, 12^ using local and public scRNA-seq data while also suggesting and substantiating a quiescent, premalignant state of primary EBV infected B-lymphocytes by contrasting/juxtaposing the primary cells with B-lymphocyte-type neoplasms, including Lymphoblastic Cell Lines (LCL).

## Results

### Viral mapping of PBMC scRNA-seq

To screen for viral transcripts, 67 publicly available scRNA-seq datasets (23.8 billion reads, corresponding to 413,902 individual cells) generated from PBMC or enriched cell types of PBMC were downloaded (Supplementary Table 1). We implemented a pipeline adapted for scRNA-seq datasets following previously described digital transcription subtraction methodology^13^. In brief, high-quality reads aligning to the human reference genome were filtered out, whereas non-human reads were retained and screened for any viral content through alignment to the NCBI’s entire collection of viral sequences. Viral reads were then manually curated and filtered for the selection of human specific viruses (Supplementary Table 1). All of the viral sequences were evaluated using Basic Local Alignment Search Tool (BLAST) analysis. This BLAST analysis revealed numerous misclassifications of either human sequences or artificial constructs as being viral reads, these sequences were omitted.

To assess the specificity of our detection method, five control datasets consisted of artificially infected primary cells (C01-C05) (Figure 1). Virus detection in the artificially EBV-infected primary B-lymphocytes (C01-C03) and HIV-infected T-lymphocytes (C04-C05) was specific to the respective agent and high levels of viral RNA corresponding to 0.04-1.07% and 0.29-0.36% of the total reads respectively was detected in these samples (Supplementary Table 1). In the samples C04 and C05, non-human viruses and virus constructs were observed in both samples, Vesicular stomatitis Indiana virus RNA was also detected in C04, all of these were considered as contaminants since the samples originated from healthy individuals.

**Figure 1.**
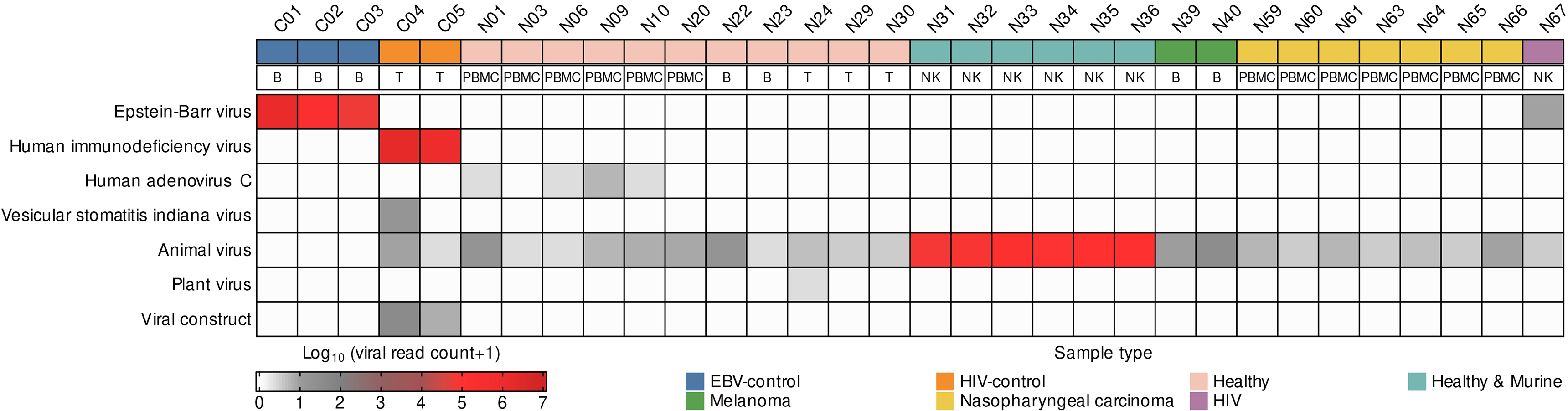
The landscape of viral sequences in public PBMC scRNA-seq datasets. Virus RNA expression levels for seven virus types and categories (vertical axis) detected in 32 datasets of scRNA-seq data from PBMCs from 17 patient cohorts (horizontal axis). Annotation in the second row denotes whether the sample is from PBMC or purified lymphocyte subpopulation.

Amongst the remaining non-manipulated datasets (N01-N67), 27 contained reads matching to the viral database. Datasets originating from healthy individuals (N01-N38) contained, in similarity to the controls C04 and C05, animal viruses. Six datasets (N31-N36) contained large amounts of murine viruses. These datasets originated from mixed human and murine NK-cells. The majority of animal virus reads from all datasets originated from murine viruses which imply that the contaminations originated from the laboratory environment. One dataset, N24, also contained plant virus RNA. Interestingly, four of the PBMC datasets (N01, N06, N09 and N10) contained human adenovirus C reads. Human adenovirus is a well-known environmental contaminant from large genomics studies, but these reads did not align to previously described regions^13, 14^ (Supplementary Figure 1). Also, the findings originate from two different studies which argue against local contamination, although plasmid vectors and cells expressing adenovirus are pervasive^15, 16^.

We continued by analyzing cancer datasets originating from B-lymphocytes of melanoma patients (N39-N40) and PBMC of nasopharyngeal carcinoma patients (N41-N50), a known EBV-associated tumor. EBV DNA is often detected in the plasma of nasopharyngeal carcinoma patients^17^. However, no EBV-infected PBMC was identified in these nasopharyngeal carcinoma datasets, confirming that the EBV DNA detected in the blood of these patients originates from the EBV-positive tumor cells either in the form of virus particles or fragments from dead cells, and not from peripherally infected lymphocytes. Contaminating animal viruses were detected in both cancer datasets, and no other viruses found. EBV or other viruses were not found in PBMC from multiple sclerosis patients (N51-N66), a disease associated with EBV^18^.

EBV DNA is often detected in plasma or PBMCs of immunocompromised patients^19^. We analyzed the NK-cells from an HIV-infected patient for viral content. Six reads mapping to the EBV genome were captured in this immunocompromised patient. These transcripts were mapped to the 3’ end of the *RPMS1* transcript, an abundantly expressed viral gene in latently infected cells^20^ (Supplementary Figure 1). Considering the tropism of EBV towards B-lymphocytes it is conceivable that the positive NK-cell is a reflection of a more prevalent B-lymphocyte infection in the same patient. Encouraged by this finding we focused on analyzing EBV expression in blood samples from immunocompromised individuals.

### EBV RNA in peripheral blood cells

Pharmacological inhibition of the cytotoxic T-lymphocyte compartment following transplantation often entails proliferation of latently EBV-infected B-lymphocytes. The severity ranges from asymptomatic viremia to fatal proliferation of EBV-transformed cells as seen in posttransplant lymphoproliferative disorder^21^. To corroborate that EBV RNA is expressed at detectable levels in circulating peripheral blood cells of immunocompromised individuals, whole-blood specimens were collected from a cohort consisting of 12 allograft recipients, ten of which have ongoing or previous history of persistent EBV viremia, and two controls. All patients were *EBNA1* seropositive, thus reflecting a state of reactivation in the setting of impaired T-lymphocyte surveillance. To determine the expression of a selection of latency transcripts (*EBER1*, *EBNA1*, *LMP-1/BNLF2a/b*, *LMP-2A* and *RPMS1*), an RT-qPCR panel with enhanced sensitivity for low abundance nucleic acids was employed as described^22^. Gene expression in relation to viral burden as determined by qPCR are presented in Table 1. Quantitative (Ct) values are provided in Supplementary Table 2.

**Table 1.** Latent EBV transcripts in peripheral blood cells. Whole-blood specimens were collected from 12 immunosuppressed allograft recipients. The samples are ordered by viral load as determined by qPCR. Levels of virus DNA are expressed as genome equivalents per milliliter (Geq/ml) blood. Expression of viral transcripts was analyzed using RT-qPCR. + symbol (gray shadow) indicates detectable levels of the respective RNA transcripts. GAPDH served as control for the RNA extraction and as a normalizer gene for relative quantification.

| Patient Data |  | EBV RNA |  |  |  |  | RNA control |
| --- | --- | --- | --- | --- | --- | --- | --- |
| Pat. No | EBV DNA<br>log Geq/ml | EBER1 | RPMS1 | EBNA1 | LMP-1/BNLF2 | LMP-2A | GAPDH |
| 1 | 4.93 | + | + | + | + | + | + |
| 2 | 4.51 | + | + | + | + | - | + |
| 3 | 4.22 | + | + | + | - | - | + |
| 4 | 3.97 | + | + | - | - | - | + |
| 5 | 3.71 | + | + | - | - | + | + |
| 6 | 3.71 | + | + | - | - | - | + |
| 7 | 3.60 | + | - | - | - | - | + |
| 8 | 3.39 | + | - | - | + | - | + |
| 9 | 2.92 | + | - | - | + | - | + |
| 10 | 2.73 | - | - | - | - | - | + |
| 11 | - | - | - | - | - | - | + |
| 12 | - | - | - | - | - | - | + |

*EBER1* was consistently detected at high levels in all individuals with ongoing viremia, with the only exception of one case with the lowest EBV-DNA concentration. As shown in Supplementary Figure 2A, the levels of *EBER1* transcripts correlated with the total viral burden in blood. Transcription from the *RPMS1* gene was detected in six out of the ten viremic individuals, more specifically in those with high viral load. The levels of *RPMS1* were moreover correlated with the levels of *EBER1* (Supplementary Figure 2B). Detectable levels of *EBNA1* transcripts were in a similar manner contingent on high viral load and restricted to the three samples with EBV-DNA levels exceeding 10,000 genome equivalents per milliliter. Detectable expression of *LMP-1/BNLF2a/b* and *LMP-2A* was on the contrary not directly correlated with high viral load. Transcription of the latent membrane proteins consequently appeared to be dispensable for cell proliferation under the specific circumstances that the cohort reflects.

Taken together, the latent transcriptional program in peripheral blood cells of viremic carriers was predominated by the short non-coding *EBER1* in particular, but also by the polyadenylated transcript *RPMS1* to a significant degree. Intriguingly, expression of the full range of latent genes was only detected in the case with the highest viral load.

### Viral gene expression in primary B-lymphocytes at single-cell level

Considering the technical constraints of scRNA-seq, including the low capture rate and limitation to polyadenylated RNA^23^, we reasoned that specific enrichment of possibly EBV-infected cells would be required for successful detection of EBV RNA. Primary B-lymphocytes were therefore isolated from a whole-blood specimen with more than five million genome equivalents of EBV-DNA per milliliter blood. *EBER in situ* hybridization (EBER-ISH) established evidence of high intracellular viral burden as a strong signal was detected in 757 cells out of 2669 total cells (rounded to approximately 30%; Figure 2A-C). The patient, a splenectomized 50 years old man with a history of Hodgkin’s lymphoma (here designated EBV patient) was admitted to hospital care with symptoms mainly consisting of fever and lymphadenopathy. Clinically, the diagnosis of primary EBV-infection was set on the basis of anti-VCA (viral capsid antigen) IgG seroconversion in paired sera in conjunction with detection of significant EBV-DNA in blood. The condition subsequently developed into a chronic course with elevated viral load persisting at markedly high levels (>100,000 genome equivalents per milliliter blood) for over 16 months.

**Figure 2.**
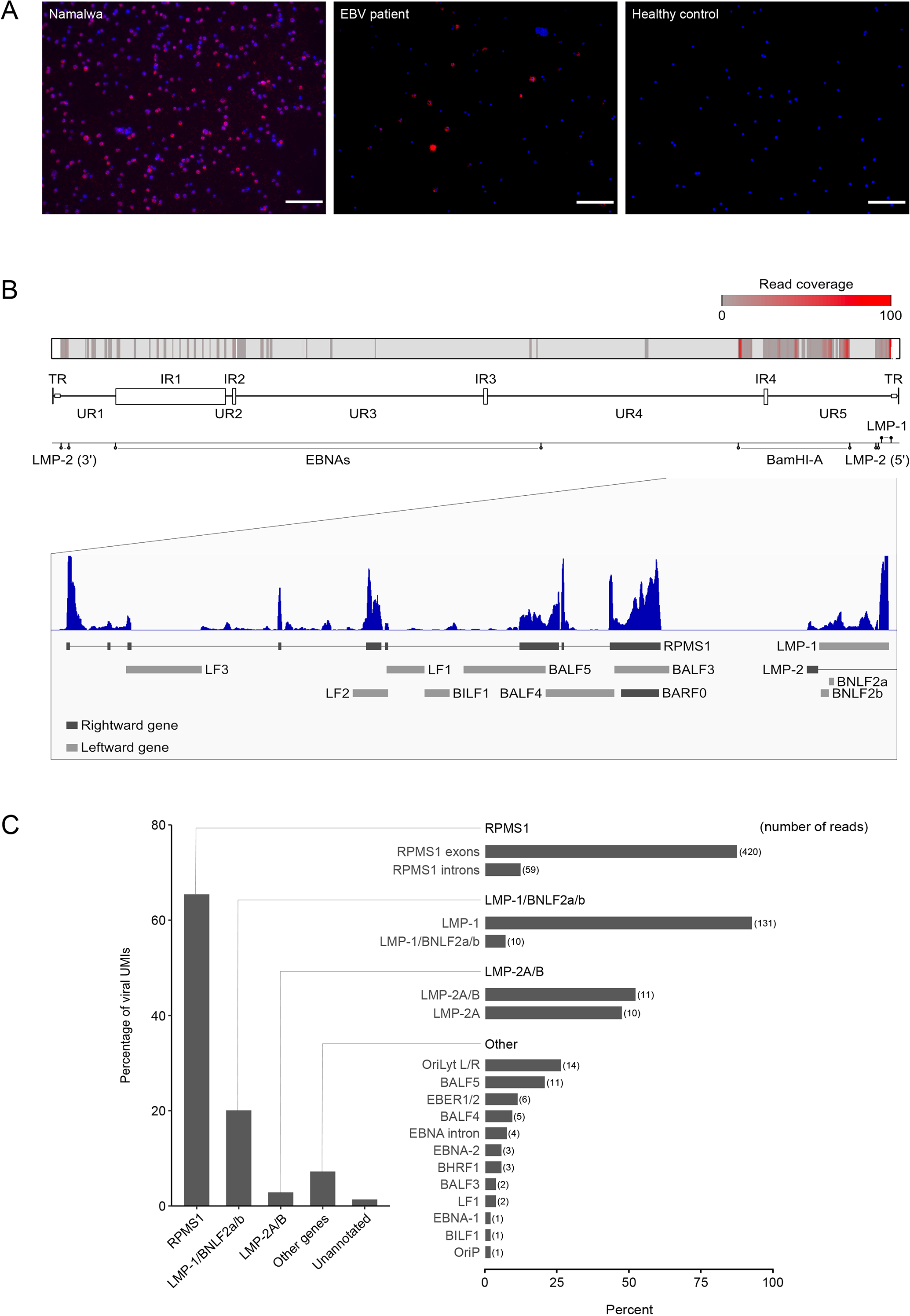
Viral gene expression in EBV infected primary B-lymphocytes. (A) Fluorescent in situ hybridization of EBER in the EBV positive Burkitt lymphoma cell line Namalwa (left) and B-lymphocytes from the EBV patient and healthy control (middle and right, respectively), scale bar: 100µm. (B) Sequencing coverage across the EBV genome NC_007605.1 with the BamHI-A and LMP-1 region highlighted. (C) Relative expression levels of EBV transcripts (right). The right diagram displays the percentage distribution within each group. Numbers in parentheses represent the absolute number of reads.

In total, 25,218 B-lymphocytes were captured with 134,403,736 unique molecular identifiers (UMIs), corresponding to 80% sequencing saturation from the EBV patient (Supplementary Table 3). Equivalent numbers were attained in the healthy control sample. In order to detect viral sequences, the EBV genome (NC_007605.1) was added as an extra chromosome to the human genome. However, only 373 UMIs were initially mapped to the EBV genome using standard settings. Here, it should be noted that Cell Ranger analysis pipelines are not adapted for analyzing compact viral genomes with overlapping genes as multi-mapped reads are discarded by default^23^. The virus reference annotation was therefore customized so as to interpret the entire EBV genome as a single gene, thereby circumventing the multi-mapping issue. In doing so, 702 viral UMIs were retrieved and 645 cells (1.09 UMI/EBV+ cells; 2.6% of total cells) were classified as EBV-positive. The low percentage of EBV-positive cells most likely reflects the limited capture rate of the scRNA-seq technology. As expected, no EBV was detected in the healthy control.

The biggest share of viral reads, corresponding to 511 reads and 73% of total EBV transcripts, were mapped to the BamHI-A and I region (NC_007605.1:138200-160500) (Figure 2B). This region contains multiple intersecting genes; however, only 25 sequences were aligned in leftward direction, indicating that the bulk of reads derived from the positive strand encoded rightward transcripts. More specifically, 420 out of the 486 rightward reads aligned to the constitutive exons of *RPMS1* (Figure 2C). Considering the abundant alternative splicing including cassette exons and intron retentions, the majority of the rightward reads which did not align to the constitutive exons most likely originated from *RPMS1* as well^24^. Thus, the overwhelming majority of viral transcripts could be ascribed to the long non-coding RNA *RPMS1*. It should yet be noted that the library preparation biasedly excluded EBERs due to the short length (<200 bp) and non-polyadenylated 3’-termini. Nevertheless, four and two reads aligned to *EBER1* and *EBER2*, respectively. EBERs could, however, be presumed to be expressed in virtually all EBV-infected cells based on the EBER-ISH and RT-qPCR results presented above.

Approximately one fifth (147 reads) of the viral reads mapped to the leftward locus in which *LMP-1* and *BNLF2a/b* are encoded. The largest part (89%) of these reads aligned upstream (5’) of the *BNLF2a/b* transcription start site in the coding region of *LMP-1*. Twenty-one reads, corresponding to 3% of total reads, mapped to the *LMP-2/BNRF1* region. These genes only share 3’ end as the majority of *BNRF1* is encoded within the introns of *LMP-2*. Five reads aligned to the overlapping *LMP-2/BNRF1* coding region, whereas the 5’ end of *BNRF1* had no coverage. These reads could therefore be assumed to derive from *LMP-2* given the 5’ library preparation. Ten reads were mapped specifically to *LMP-2A* while eleven reads were aligned to the shared *LMP-2A/2B* exon.

No other viral transcripts exhibited any coverage equivalent to more than 2% of total viral sequencing reads. As shown in Figure 2C, 46 reads could be assigned to various positions in the genome, e.g. *OriLyt L/R*, *BALF5*, *BALF4* and *EBNA* intron segments. In conclusion, the single-cell EBV transcriptome analysis clearly highlighted the ubiquitous expression of *RPMS1*. However, considering that only one viral UMI was detected in the vast majority of EBV-positive cells, the scRNA-seq data reflects the quantitatively most representative transcripts and does not rule out concomitant expression of other genes. Nonetheless, the high expression of *RPMS1* observed in this patient is in line with the RT-qPCR results.

### B-cell receptor sequencing of EBV-infected lymphocytes

Insight into the peripheral B-cell receptor repertoire at single-cell level allows for highly sensitive detection of clonality and, moreover, linkage of EBV to specific points in the B-lymphocyte development based on the distinctive immunoglobulin gene rearrangements and somatic hypermutations for each clonotype. Neither the EBV patient nor the healthy control displayed any apparent monoclonal cell populations as determined by the Gaussian distribution of the length of the third complementary region (CDR3) of the heavy chain (IGH) (Supplementary Figure 3). As regards to the immunoglobulin variable (V) region genes, elevated usage of IGKV3-20 and IGHV1-69 was observed in the EBV patient (Supplementary Figure 4). A preferential usage of these particular V genes has previously been noted in Burkitt’s lymphoma and multiple sclerosis^25, 26^. However, IGKV3-20 was also overrepresented in the healthy control.

The CDR3 sequences were further used to assign a unique identity to each clonotype and thereby determine the total cell count for each clone. The number of clonal cells in relation to total cells, here denoted clonal expansion factor, was higher in the patient compared to the healthy control, which pointed to a subtle polyclonal proliferation. Indeed, the frequency of relatively large clonal populations was significantly higher in the patient compared to the healthy control. (Figure 3A). As might be expected, EBV-positive cells were accordingly more likely to be clonally expanded compared to EBV-negative cells (Figure 3B). Eight out of the ten largest B-lymphocyte clones turned out to contain at least one EBV-infected cell; however, this number could presumably be higher due to the low capture rate.The polyclonal outgrowth in the patient was consequently most probably driven by EBV, as such B-lymphocyte expansions did not exist in the healthy control. All of the ten largest clonal populations exhibited evidence of somatic hypermutation and class-switch recombination in the IGH loci (Supplementary Table 4). This indicated a history of passage through a germinal center-like reaction prior to entering the peripheral compartment. It is also worth noticing that no significant intraclonal sequence variation was observed within expanding clonotypes, which argues against ongoing somatic hypermutation. Furthermore, the number of false EBV-negative cells could be extrapolated based on the presumption that all cells with shared clonal origin should be considered as EBV-positive if merely one cell within that given clonotype was classified as such. In doing so, the number of EBV-infected cells was estimated to 1,657 cells (6.6%).

**Figure 3.**
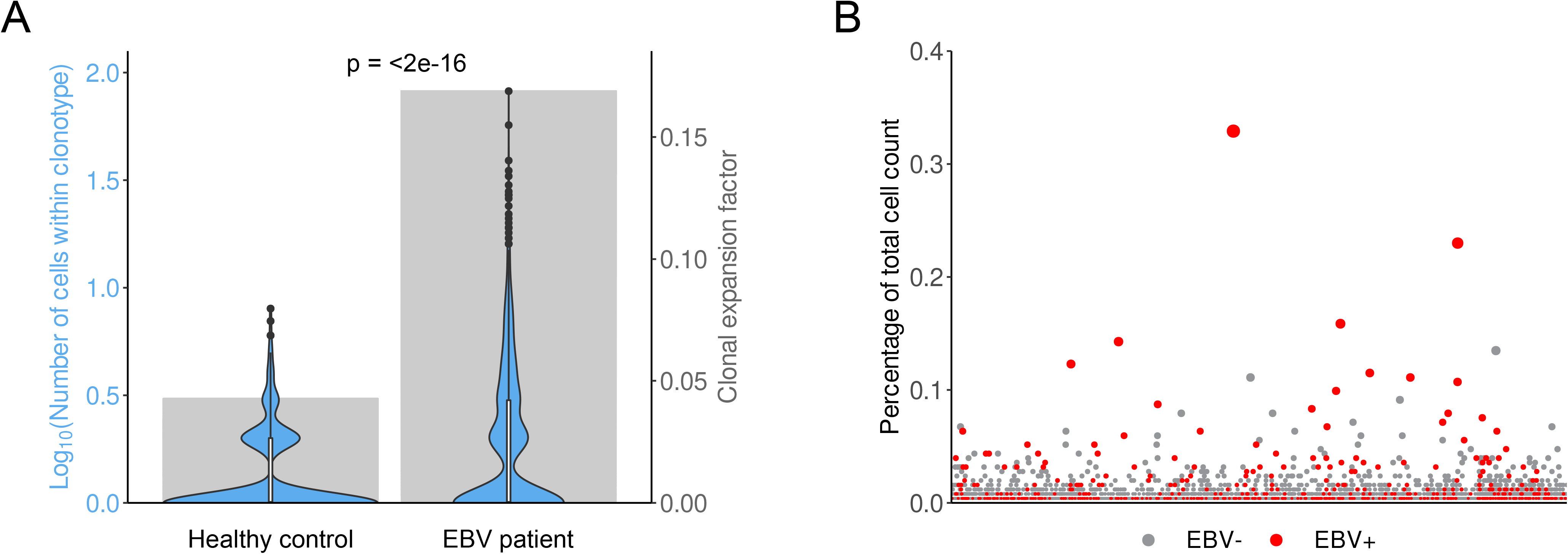
Clonal expansion as determined by the B-cell receptor. (A) Distribution of number of cells within each clonotype (violin plot, left y-axis) and number of clonal cells in relation to the total number of cells (gray bar plot, right y-axis). The number of cells per clonotype is significantly higher in the EBV patient compared to the control (Wilcoxon rank-sum test, p < 2.2e-16). (B) Size of each clone expressed as the percentage of total cell count. Red color indicates that EBV transcripts are detected in at least one cell.

### Clustering and Differential Gene Expression

To identify in which B-lymphocyte subpopulation the EBV-infected cells reside and characterize the transcriptome perturbations caused by the EBV-transformation we performed an unbiased clustering of the datasets, Uniform Manifold Approximation and Projection for Dimension Reduction (UMAP; Figure 4, Supplementary Figure 5). Two distinctive clusters were observed in both samples and as expected one cluster almost exclusively expressed *CD27*, denoting the memory B-lymphocyte population (right clusters). This CD27+ population could further be divided into IgM/IgD+ or IgM/IgD-subclusters. The opposite major clusters contained the CD27-IgM/IgD+ naïve B-lymphocyte population (left clusters). The absolute majority of the EBV-infected cells (99.6%) were found within the CD27+ memory B-lymphocytes clusters as expected^27^.

**Figure 4.**
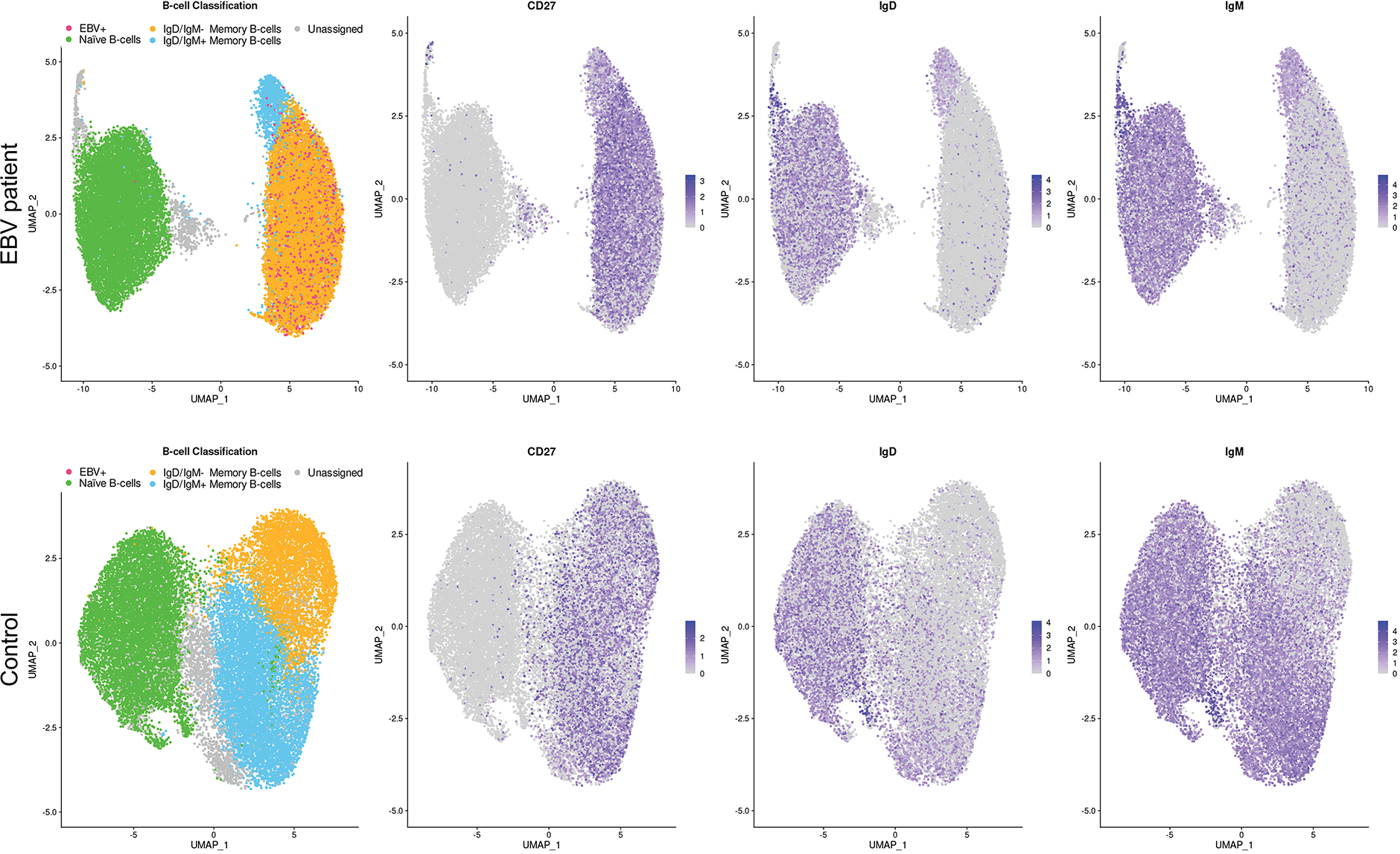
Single-cell analysis of B lymphocytes from the EBV patient and the control. Two distinct subpopulations formed in UMAP clustering of B-lymphocytes from the EBV patient (upper row) and the control (bottom row). Naïve (left cluster, orange) and mature (right, green and blue) B-lymphocytes (first column) displayed distinct expression of CD27 (second column), IgM (third column) and IgD (fourth column). EBV-positive cells are marked in red.

To elucidate the difference between EBV-positive and EBV-negative memory B-lymphocytes in the EBV patient in terms of host cell expression pattern we categorized the EBV-positive cells, explicit or inferred by clonotype assignment, into their own cluster. Due to the small number of CD27+ IgM/IgD+ lymphocytes we continued to only consider the CD27+ IgM/IgD-population. A differential gene expression analysis between these assigned EBV-positive and EBV-negative CD27+ IgM/IgD-B-lymphocytes resulted in only two genes being significantly differentially, *DUSP2* and *MYADM*, besides EBV genes. Considering the discrepancy between the fraction of EBV-positive cells detected in EBER-ISH and scRNA-seq, as well as taking into account the technical limitation of scRNA-seq, it is reasonable to assume that a large number of EBV-infected cells were assigned to the EBV-negative category which dilutes the differences between the groups.

In order to circumvent this problem we merged and integrated our two datasets so as to compare the inferred EBV-infected cells with a true EBV-negative memory B-lymphocyte population, designated local integration (Figure 5A, left and middle panel). A parallel integration was performed using three publicly available B-lymphocyte single-cell datasets (N21-N23) in order to address concerns about differing RNA expression profiles based upon individual base-level differences rather than based on the presence or absence of EBV, designated public integration (Figure 5A, right panel). Initial estimation of integration performance within both the local and public integration show a uniform mixture throughout each respective UMAP. As expected within the local integration, the major divide between the cells are between the memory B-lymphocytes (yellow clusters) and the naïve B-lymphocytes (green clusters) classified by expression levels of CD27, IgD, and IgM (Supplementary Figure 6). In the local integration, the naïve B-lymphocytes and the IgD/IgM-memory B-lymphocytes were subset into separate datasets. The differential gene expression comparing the EBV patient naïve B-lymphocytes to the control naïve B-lymphocytes serves as a control for host-specific differences and general humoral factors which would affect the naïve and memory B-lymphocyte populations. The overlapping genes from the analysis of the naïve B-lymphocytes with the analysis of inferred EBV+ memory B-lymphocytes compared to the control memory B-lymphocytes would therefore be non-significant. Using heuristic thresholds, 84 and 142 genes showed significant levels of differing expression in the naïve B-lymphocytes and memory B-lymphocytes, respectively (Figure 5B, Supplementary Table 5). In the public integration, the CD27+ IgD/IgM-memory B-lymphocytes were subset into a separate dataset so as to perform a differential gene expression comparing each one original dataset (N21, N22, and N23) with the other two datasets. This comparison generated a list of 114 uniquely and significantly differentially expressed genes between three individuals which would further be used as a separate control for finding host-specific differences (Supplementary Table 5). When using both the differentially expressed genes in the naïve B-lymphocytes and the public integration as an intermediary to filter out possible extraneous determinants, more specifically variation in gene expression due to individual host-specific differences, 97 genes remained (Figure 5C). Using this curated list of 97 differentially expressed genes, the genes were separated based on their direction of change (i.e increased/decreased). While the number of downregulated genes (n = 26) were insufficient to generate a statistically significant enriched pathway, the upregulated genes (n = 71) were analyzed for biologically functional pathway enrichment which showed an enrichment in 28 pathways (Figure 5D). Interestingly, the pathway which showed highest statistical significance was “regulation of cell population proliferation”, suggesting that the presence of EBV in memory B-lymphocytes upregulates proliferation, which is supported by the observation of polyclonal expansion of EBV-positive clonotypes (Figure 3). Additionally, three of the top seven pathways with highest statistical significance was “negative regulation of interleukin-2 production”, “cellular response to tumor necrosis factor”, and “cellular response to interferon-gamma”, implying a transcriptomal perturbation conducive towards evading the immune system. However, the relatively subtle transcriptomic effects imply that these cells represent a selected population of quiescent EBV-infected B-lymphocytes, as cells that undergo uncontrolled proliferation and/or reactivation may elicit an immune response and be eradicated.

**Figure 5.**
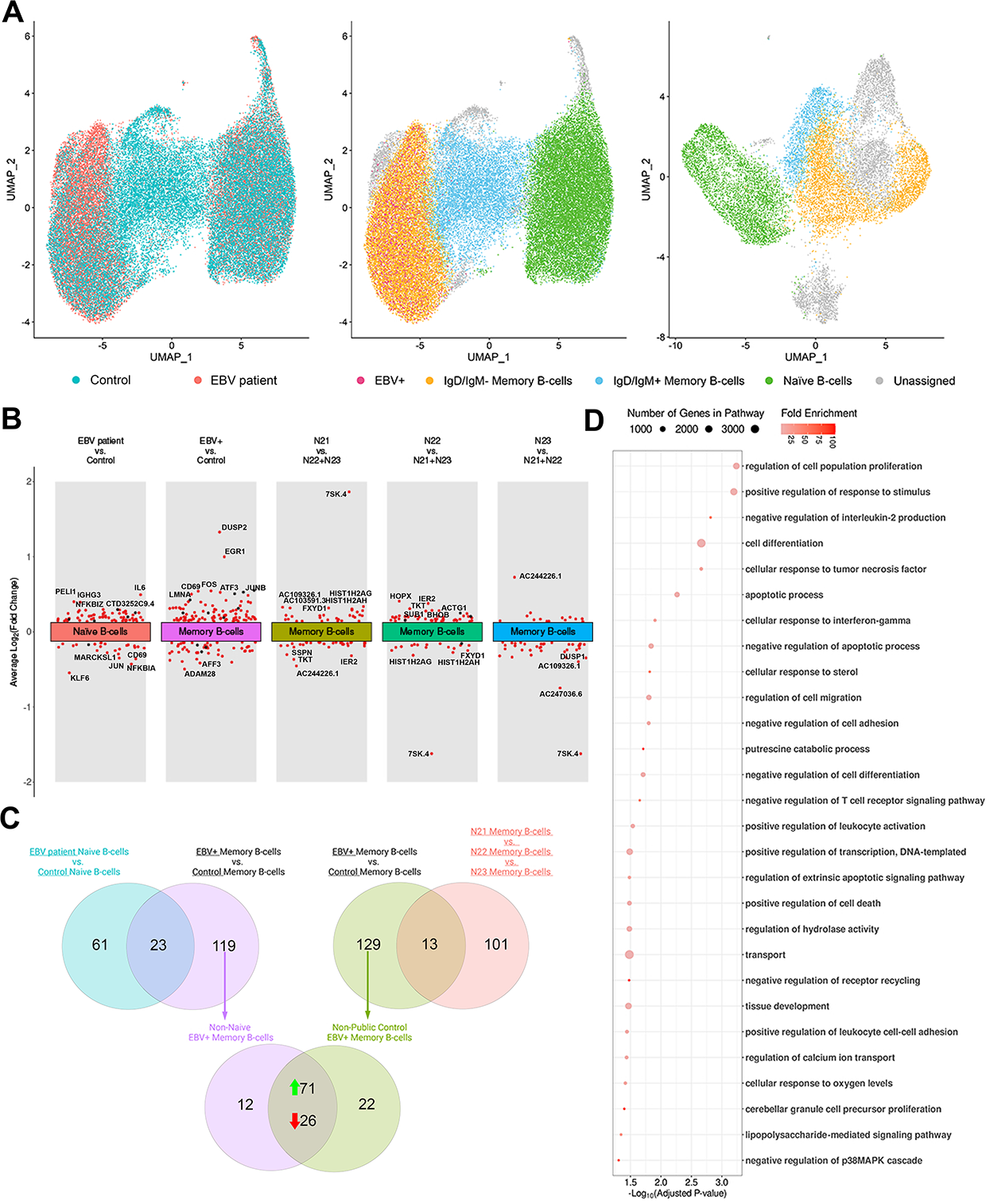
Integrated single-cell analysis and differential gene expression analysis of EBV patient filtered to four control datasets. Single-cell and differential gene expression analysis of the EBV patient and the four controls. (A) UMAP visalizations of the B-lymphocytes highlighting naïve B-lymphocytes (middle and right panel, green) and mature B-lymphocytes (middle and right panel, yellow). EBV positive cells are highlighted in red. Additionally, the performance of the integration between the EBV patient and the control, I.e the local integration, is depicted (left panel). (B) Volcano plot visualizing five different differential gene expression analysis with the y-axis representing the average log2FC change, red dots representing genes that have an adjusted p-value below 0.01, and dots annotated with gene names representing each comparisons highest differentially expressed gene. (C) Filtering of individual host-specific differences visualized through Venn-diagrams with 71 upregulated genes and 26 downregulated genes. (D) Gene Ontology Pathway Enrichment analysis of the 71 upregulated genes with the x-axis representing −log_10_(Adjusted P-value).

### Primary EBV-infected B-lymphocytes as potential cell models

EBV B-lymphocyte cell models are either derived from primary lymphoma or *in vitro* transformation of primary B-lymphocytes into Lymphoblastoid Cell Lines (LCL). Cell lines derived from lymphoma patients have a propensity to lose its EBV genome during passage of the cells. In contrast, high expression of EBV genes is readily observed in LCLs generated in the absence of immunological surveillance. To examine the difference between primary EBV-infected B-lymphocytes and cell models we merged and integrated; the EBV patient, the control, the public controls (N21-N23), the LCLs (C01-C03), and a collection of four diffuse large B-cell lymphomas (DLBCL). As there are currently no scRNA-seq available for EBV-positive B-lymphocyte malignancies we included the DLBCL to represent neoplastic cells. In total, 80,957 cells were merged and clustered into 18 different clusters (Figure 6A).

**Figure 6.**
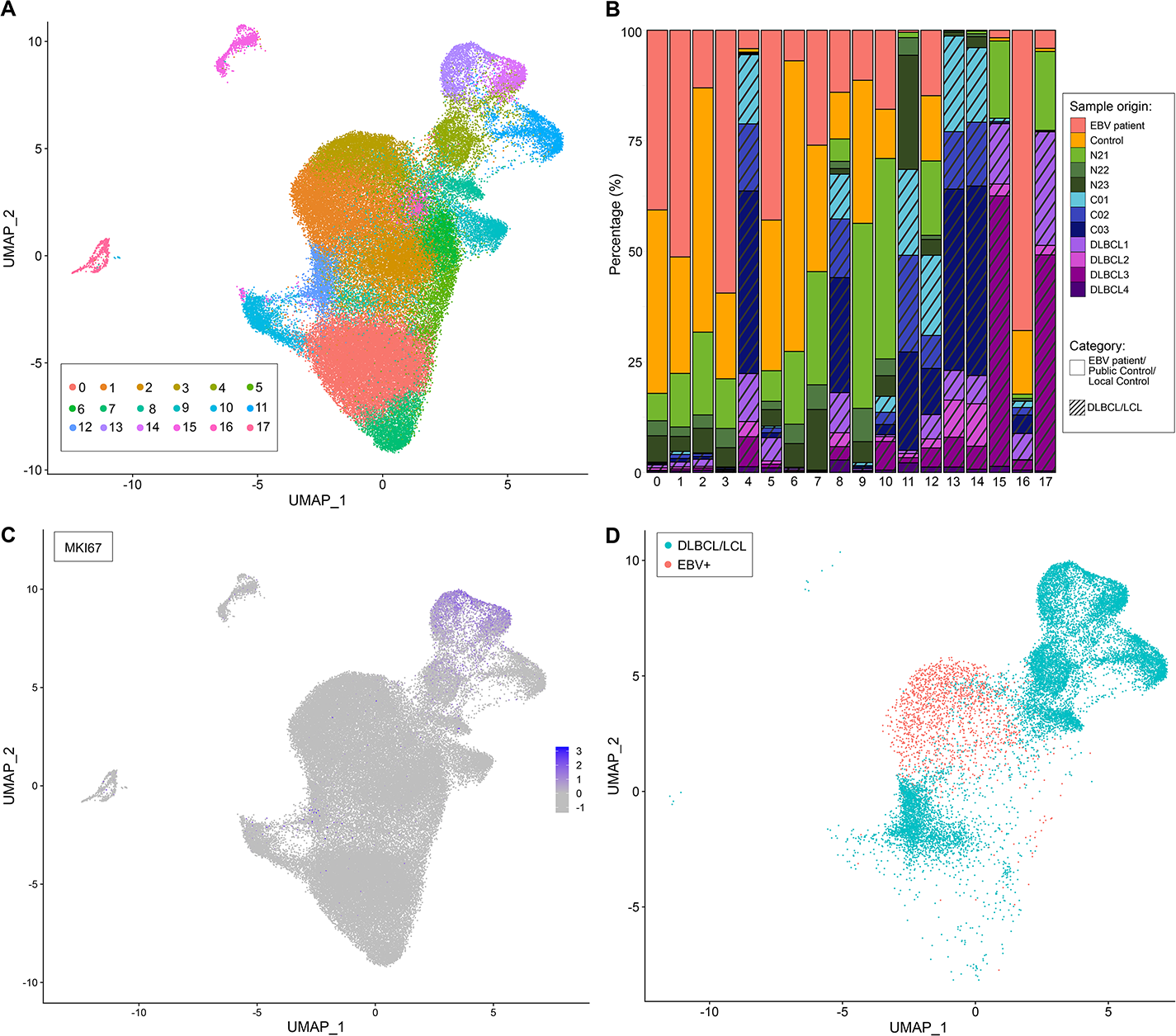
Integrated single-cell analysis of the EBV patient, four healthy controls, three LCLs, and four DLBCLs. Single-cell analysis of the integration between the EBV patient, four healthy controls, three LCLs, and four DLBCLs highlighting the malignant cells. (A) UMAP visualization and unbiased clustering of the 12 datasets, representing 80957 individual cells, split into 18 different clusters. (B) Barplot depicting the dataset composition of the 18 clusters with stripes denoting malignant origin. Red represents the EBV patient; orange, the control; shades of green, the three public controls; shades of blue, the three LCL samples; and shades of purple, the four DLBCL samples. (C) Feature plot depicting the relative expression of the cellular proliferation marker MKI67. (D) UMAP visualization when subsetting the EBV-positive B-lymphocytes as well as the cells from cluster with more than 50% cells originating from any of the malignant datasets (DLBCL/LCL)

The EBV patient, control, and public controls formed as previously clusters with naïve (clusters 0 and 7) and mature (cluster 1, 2, 3 and 6) B-lymphocytes (Figure 6B). These datasets contain the very low proportions of the cancer cells from DLBCL and the in vitro EBV-transformed LCL. Clusters 15 and 17 were clearly separated from the other clusters and the majority of the cells originated from the DLBCL datasets. The gene expression profile of these clusters suggest that they are tumor infiltrating lymphocytes (TIL) in which cluster 15 has the expression profile most similar to NK/T-lymphocytes while cluster 17 has the expression profile most similar to monocytes (Supplementary Figure 7). This clear separation of cluster 15 and 17 apart from the B-lymphocytes and simultaneous proximity to the DLBCL and the LCL B-lymphocytes suggests that the initial integration removed large parts of the batch effect while retaining the biologically relevant variance.

The DLBCL and LCL cells made up more than 50% of the cells in clusters 4, 8, 11, 13 and 14. These clusters displayed an increased expression of MKI67 with cluster 4, 13, and 14 showing an average log2 fold change above 0.10 implying that these are the clusters where the proliferative cells aggregate (Supplementary Table 6). This is further supported by the UMAP visualization of the relative expression of MKI67 for each cell where cluster 13 and 14 show clear distinction in comparison with the other clusters (Figure 6C). The inferred primary EBV-infected cells from the EBV patient remains evenly dispersed in the clusters designated IgD/IgM-memory B-lymphocytes (Supplementary Figure 8, Supplementary Figure 9) and does not cluster to the DLBCL or LCL cells (Figure 6D) to a large extent. The lack of clustering of LCL with the primary EBV-infected B-lymphocytes shows that the transcriptomic similarities may be more dependent on cell state rather than EBV gene effect.

The discrepancy between the LCL and DLBCL datasets and the controls and EBV patient can further be observed in the total number of transcripts per cell quantified by the total number of UMIs per cell in correlation with its sequencing saturation (Figure 7). In the LCL datasets (C01-C03), an increasing number of EBV genes detected is correlated with an increased number of RNA (UMIs), with the exception of the cells with most EBV genes in C01 and C02 (yellow cells bottom right). UMAP-clustering of the LCL datasets showed a distinct population in C01 and C02, but not C03 (Supplementary Figure 10). These subclusters in C01 and C02 had a high number of EBV genes expressed coupled with a low number of total (host and EBV) mRNAs in the cell. These cells with abundant EBV gene species most likely represent reactivating cells with a host shut-off. The public controls, N21-N23, and the EBV patient have a sequencing saturation spanning from 77-87% and a small minority of cells contain more than 2,000 mRNAs per cell (0.04-1.50%). In contrast, a large proportion of the cells in the LCLs contained more than 2,000 mRNAs (UMIs) per cell (26.9-36.3%), despite only having a sequencing saturation of 31-47%, a similar observation has previously been made in bulk sequencing datasets^28^. An intermediate level of RNA expression was observed in the DLBCL datasets with an equivalent sequencing saturation in comparison with the controls. Thus the primary EBV-infected B-lymphocytes represents a quiescent premalignant EBV-transformation with a subtle increased proliferation and a retained sparse RNA expression and may be appropriate as a cell model for the studies of viral induced host transcriptome perturbation.

**Figure 7.**
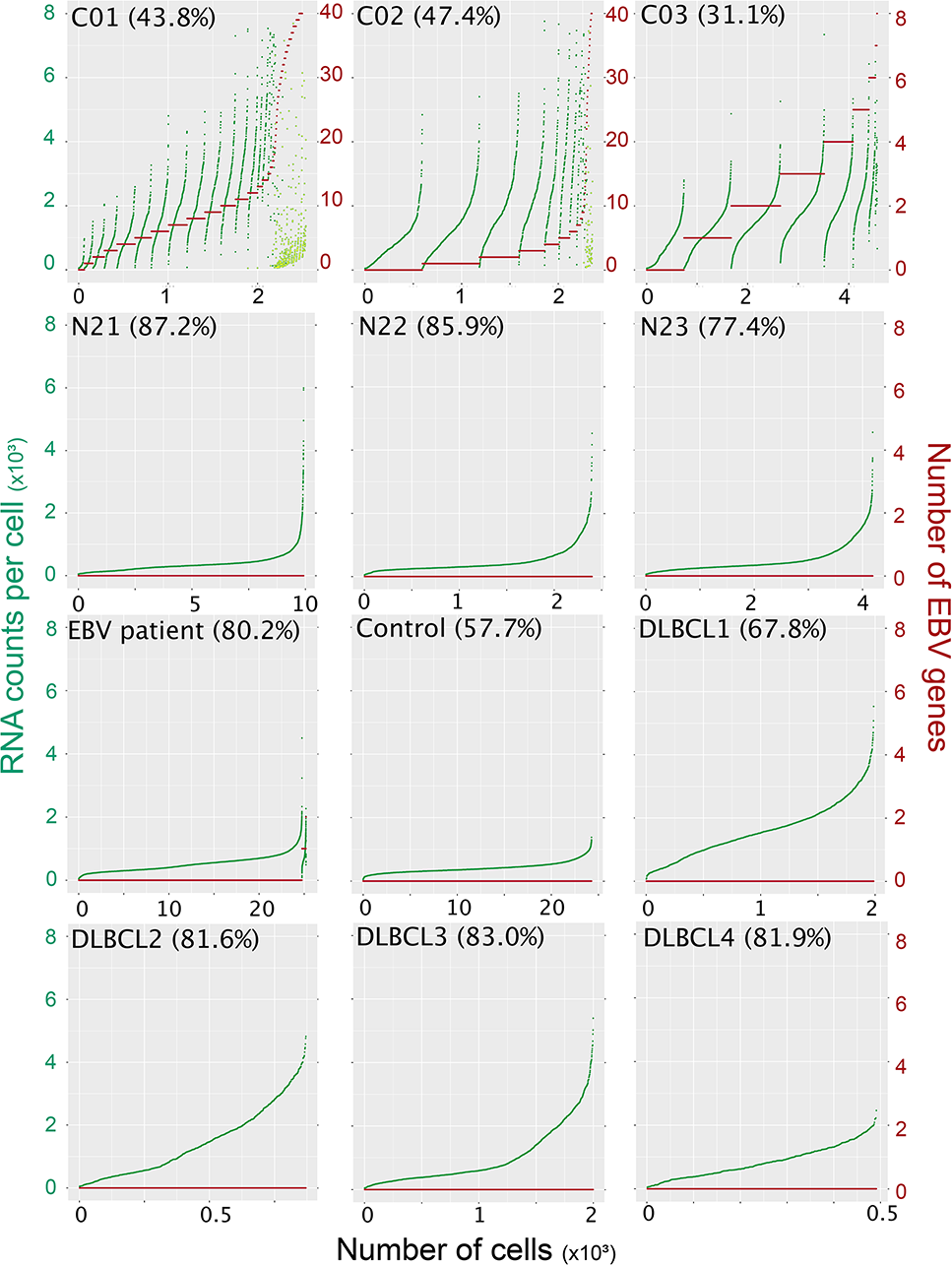
Distribution of number of EBV genes and RNA content per cell. The total number of UMI’s per cell (green dots, left y-axis) is plotted in increasing order categorized by the number of EBV genes per cell and total RNA count per cell (red, x-axis). Differentially clustered subpopulations in LCL-datasets (top row, yellow dots) C01 and C02 displayed high numbers of EBV genes (different left y-axis scale) and low RNA content representative of lytic cells with host shut-off. No EBV RNA was detected in controls (second row and Control) or the DLBCL datasets. Host RNA expression in EBV-positive cells in the EBV patient displayed a comparable level compared with EBV-negative cells (third row left). Sequencing saturation is indicated in parenthesis for each dataset.

## Discussion

Primary latently EBV-infected B-lymphocytes in peripheral blood are sparse and the inability to enrich these cells has impaired the understanding of EBV-transformation in non-cancerous cells. We have in this study identified that patients with an inhibited immune system have EBV RNA expressing cells in the peripheral blood at sufficient quantities for detection using enhanced sensitivity RT-qPCR and single-cell analysis. Bioinformatics adaptation using a modified virus reference genome and coupling of VDJ sequencing data further increased the classification of EBV-positive cells almost five-fold to encompass 6.5% of all B-lymphocytes in scRNA-seq. However, this fraction is 5-fold lower than the results obtained from EBER-ISH. Thus, the main obstacle for viral read detection using scRNA-seq lies in the capture rate and the polyA-capture used for creating cDNA. The highly expressed non-polyadenylated viral transcripts EBER and for other herpesviruses latency-associated transcripts are not compatible with the UMI-tagged polyT-primers for the generation of a global cDNA library for scRNA-seq. Nevertheless, these limitations produce a high proportion of false negative cells and thus impairs a proper comparison of virus-infected and non-infected cells, especially if the fraction of virus-infected cells are high and is coupled with a low expression of polyadenylated viral RNA. For future studies, an increased sensitivity for scRNA-seq could be achieved using methodology with higher capture rate or a complementary strategy to utilize targeted EBV genes, including non-polyadenylated RNA, into the capture probes or enrichment of EBV-positive cells prior to library building.

In accordance with previous findings we find in the scRNA-seq data that EBV is harbored in the CD27+ B-lymphocytes. We could further identify polyclonal outgrowth of certain clonotypes. Importantly, despite the EBV-infected B-lymphocytes co-clustered with mature B-lymphocytes from healthy individual, i.e. showing high similarity with normal B-lymphocytes, an induction of pro-proliferation (and immune evasion) genes were observed in the EBV-infected cells. This propensity for proliferation coupled with a low EBV and host gene expression can be put in contrast with the highly proliferating LCL which expresses a large number of EBV and host genes that could elicit an immune response. Furthermore, enriched genes in the EBV-infected B-lymphocytes e.g. CD80 could provide markers for further enrichment of primary latently infected cells.

Our workflow consisting of patient category selection, enhanced sensitivity RT-qPCR, *in situ* hybridization and cell type enrichment as well as bioinformatic pipelines provides a comprehensive guide for scRNA-seq of other virus-infected cells in blood. The scRNA-seq methodology presented here can be used for functional and descriptive studies of EBV-transformation and maturation process of B-lymphocytes by longitudinal sampling of patients prior to and after transplantation/infection. The expansion and shrinkage of different EBV-infected B-lymphocyte subpopulations, the cells’ proliferative capacity and the virus propensity to reactivate could be characterized at the cellular level in conjunction with the immunological pressure. Transformation from a premalignant quiescent infection into a malignant neoplasm during acquisition of somatic mutations would be possible to study in individuals with high risk for developing post-transplant lymphoproliferative disease. Lastly, identification of the EBV-infected reservoir would allow for eradication of the latent virus *ex vivo* in bone marrow used for transplantation.

Our results exemplify that EBV-infected primary cells are thus suitable for studies of EBV programming and the cellular effects of the expressed EBV elements in quiescent cells with the ability to avoid the immune system, and in the absence of somatic mutations driving proliferation.

## Supplementary information

Supplementary Table 1: General sequencing details in PBMC scRNA-seq dataset

Supplementary Table 2: Ct values of EBV transcripts in the 12 samples and primers for (RT-)qPCR

Supplementary Table 3: Summary of in-house datasets using different EBV references

Supplementary Table 4: Somatic recombination and hypermutation of the BCR

Supplementary Table 5: Differential Gene Expression in Figure 5

Supplementary Table 6: Differential Gene Expression in Figure 6

Supplementary Figure 1: Aligned viral reads in scRNA-seq datasets

Supplementary Figure 2: Correlation between viral load and EBV transcripts

Supplementary Figure 3: CDR3 length distribution

Supplementary Figure 4: Immunoglobulin variable region gene usage

Supplementary Figure 5: Initial clustering of the EBV patient and control

Supplementary Figure 6: Initial clustering of the integration between EBV patient and control

Supplementary Figure 7: Cell-type marker expression

Supplementary Figure 8: Manual relabeling of unbiased clustering in the integration between EBV patient, four controls, three LCLs, and four DLBCLs

Supplementary Figure 9: B-lymphocyte subtype markers

Supplementary Figure 1. Aligned viral reads in scRNA-seq datasets

(A) Overview of reads aligning to the Human adenovirus C genome(top panel). Detailed view of reads aligning to E1A (i), E1B (ii) and E1B/IX (iii). (B) Overview of reads aligning to the Epstein-Barr virus. Detailed view of reads aligning to RPMS1 (i).

Supplementary Figure 2. Correlation between viral load and EBV transcripts

(A) Correlation between EBV DNA levels quantified by qPCR and transcription of EBER1. EBV transcripts were measured using targeted preamplification followed by RT-qPCR and values on the x-axis correspond to the difference in Ct values between the housekeeping gene *GAPDH* (without preamplification) and EBER1. Higher levels of EBER1 correlates with an increase in viral load (Pearson R^2^ = 0.83, p = 0.00068). (B) The relative expression levels of EBER1 and RPMS1 were correlated (Pearson R^2^ = 0.86, p = 0.0083).

Supplementary Figure 3. CDR3 length distribution

Length distribution of the immunoglobulin heavy chain complementarity-determining region 3 (CDR3) with a Gaussian distribution curve in both the EBV patient and healthy control.

Supplementary Figure 4. Immunoglobulin variable region gene usage

Immunoglobulin heavy and light chain variable region gene usage in the healthy control and EBV patient.

Supplementary Figure 5. Initial clustering of the EBV patient and control

Figures displaying the EBV patient’s (upper) and the control’s (lower) UMAP visualization and expression of B-lymphocyte subpopulation marker genes (CD27, IGHD, and IGHM) as dotplots.

Supplementary Figure 6. Initial clustering of the integration between EBV patient and control

UMAP visualization, unbiased clustering, each clusters relative expression of cell-type markers in violin plot format and dot plot format of the integrated scRNA-seq data between EBV patient and the control.

Supplementary Figure 7. Cell-type marker expression

Each clusters relative expression of cell-type markers in violin plot format and dot plot format of the integrated scRNA-seq data between EBV patient, control, public control, LCL, and DLBCL.

Supplementary Figure 8. Manual relabeling of unbiased clustering in the integration between EBV patient, four controls, three LCLs, and four DLBCLs

UMAP visualization of the integrated scRNA-seq data between EBV patient, control, public control, LCL, and DLBCL after relabeling all 18 clusters into one of eight classes: IgD/IgM-Memory B-cells, IgD/IgM+ Memory B-cells, Naïve B-cells, DN Memory B-cells, NK/T-cells, Monocytes, DLBCL/LCL, and Unassigned.

Supplementary Figure 9. B-lymphocyte subtype markers

FeaturePlot visualizing each individual cell’s relative expression of typical B-lymphocyte subtype markers (CD27, IgD, and IgM) as well as the cells relative expression of PCNA as an additional marker for cellular proliferation.

Supplementary Figure 10. The B-lymphocyte subtype clusters in LCLs

UMAP visualization of an unbiased clustering of the three samples of LCL. Clear subclustering was visible in C01 and C02 with cluster 2 and cluster 10 having a higher proportion of EBV transcripts to total RNA, respectively. Cluster 2 in C01 and cluster 10 in C02 are indicative of reactivating cells. C03 does not seem to show any apparent subclustering and subsequent population of EBV reactivating cells.

Supplementary Table 1. General sequencing details in PBMC scRNA-seq datasets

Viral screening of publicly available scRNA-seq data from 16 studies equivalent to five control datasets and 67 individual datasets. Each dataset’s metadata as well as raw number of viral sequences are reported. The number of each dataset’s virus aligning sequences are also depicted in the table. Identical report was done for the scRNA-seq data from the EBV patient and the control.

Supplementary Table 2. Ct values of EBV transcripts in the 12 samples and primers for (RT-)qPCR

Supplementary Table 3. Summary of in-house datasets using different EBV references

Supplementary Table 4. Somatic recombination and hypermutation of the BCR Genetic features/characteristics of the B-cell receptor genes in the most expanded clones in the EBV patient.

Supplementary Table 5 - Differential gene expression between EBV patient and the control as well as between the public controls

Differential gene expression analysis from three comparisons using EBV patient, control, and the public controls.

Supplementary Table 6 - Differential gene expression between the integration of EBV patient, control, public control, DLBCL, and LCL

Comprehensive differential gene expression analysis from the integration between EBV patient, control, public control, DLBCL, and LCL. Unbiased clustering was performed wherein the differential gene expression compared the expression of one cluster against all other clusters.

## Materials & Methods

### Viral detection in PBMC scRNA-seq

Single-cell RNA sequencing data in bam and fastq-format was obtained from a web-based repository. Datasets originating from 10x Genomics own repository were downloaded using the wget function. Datasets originating from EMBL-EBI ArrayExpress repository were downloaded using the wget function in fastq-format. All other datasets were downloaded using sratools (version 2.9.6) in bam or fastq format. Samples in bam format were processed into fastq format using Cellranger’s (Version 3.0.1) function bamtofastq.

All processed fastq files had a Unique Molecular Identifier (UMI) sequence and a Cellular Barcode (CB) attached to the reads with UMI-tools (Version 1.0.0). Low-complexity reads were filtered and removed using PRINSEQ (Version 0.20.4). Next, the reads were sorted based on affinity for aligning with the human genome using Bowtie Aligner with the GRCh38 reference. All non-aligned reads were further processed. The non-human reads were aligned with Bowtie Aligner to a collection of viral genomes from NCBI Reference Sequence (RefSeq). The specific parameters are as follows: For prinseq: Minimum read length of 45 nucleotides, minimum read quality of 25, low complexity sequence filter method “Dust” with a threshold value of 7. For Bowtie human read filtering: While using the GRCh38 as reference genome, maximum number of misalignments of 2, maximum number of valid alignments per read/pair of 1 and aligned read output to discard (`/dev/null/`). For Bowtie viral sequence detection: While using NCBI’s RefSeq collection of viral genomes (n=3,590) as reference, maximum number of misalignments of 2, maximum number of valid alignments per read/pair of 1, reported singleton alignments are the highest quality in terms of stratum of “`TRUE`”, and maximum number of reportable alignments of 25. The viral aligned fastq files, in bam file format, were then processed with an in-house Python script to extract and count the number of viral reads. The output was formatted into a data matrix according to the tab-separated value standard and the resulting data matrix file was used to represent the viral content of each dataset. For subsequent refined analysis, datasets that contained zero viral reads were not processed further. Datasets from artificially infected cells (C01-C05) were also omitted. Multiple datasets (N31-N36), containing mouse-specific viruses, were confirmed to contain mouse cells through alignment to the mouse reference genome (GRCm38) using Bowtie Alignment and were also excluded from further analysis. The remaining datasets were analyzed with the Basic Local Alignment Search Tool (BLAST) to verify that the pipeline and alignments were correct.

### EBV quantification of immunocompromised cohort

Blood samples submitted for EBV DNA quantitative PCR at the Department of Clinical Microbiology at Sahlgrenska University Hospital during a one-week period were collected. Transplanted patients with previous serological results indicating EBNA IgG reactivity and PCR positive values were further sorted and twelve samples representing a range of EBV DNA load were selected. Nucleic acid extraction was performed using the MagNA Pure Compact DNA Isolation Kit I (Roche Diagnostics) on the MagNA Pure LC automated extractor. An input of 200 µl whole-blood spiked with phocine herpesvirus 1 was eluted to a 100 µl extraction buffer. EBV DNA quantitative real-time PCR was run as originally described^29^, with modifications previously reported^30^.

### RNA extraction and reverse transcription

Total RNA was extracted from 100 µl of whole-blood specimens (diluted in 150 µl PBS) using TRIzol LS Reagent (Invitrogen/Life Technologies/Thermo) according to the manufacturer’s instructions. Reverse transcription was performed with TATAA GrandScript cDNA FreePrime Kit (TATAA Biocenter). A reaction containing 5 µl (200 ng - 500 ng) RNA, 1x TATAA GrandScript RT Reaction Mix, 1x TATAA GrandScript RT enzyme, random hexamers and oligo(dT)20 primers were added to a final volume of 20 µl and incubated at 25°C for 10 min, 42°C for 45 min, 85 °C for 5 min and then held at 4°C.

### Targeted preamplification and quantitative real-time PCR

Multiplex target-specific preamplification was performed as described^22^. The same set of primer pairs were used for preamplification and subsequent RT-qPCR (sequences are provided in Supplementary Table 2). Two microliter of cDNA was preamplified in a 10 µl reaction containing 2X TATAA SYBR GrandMaster Mix (TATAA Biocenter), 400 nM of each primer pair, 1 µl of 10 mg/ml bovine serum albumin and 1 µl 25% glycerol. Initial heating at 95°C for 3 min was followed by 23 cycles of amplification (95°C for 20 s, 60°C for 3 min and 72°C for 20 s) and 10 min of final elongation at 72°C. Subsequent quantitative PCR was performed using SYBR GrandMaster Mix (TATAA Biocenter). Two microliters of preamplified template diluted 1:20 in TE buffer were transferred into a 20 µl reaction mixture containing 2X TATAA SYBR GrandMaster Mix and 0.5 uM primers. The following thermal protocol was applied: 95°C for 2 min prior to 40 cycles of amplification (95°C for 5 s, 60°C for 20 s and 72°C for 20 s). Reaction specificity was evaluated by melting curve analysis with the dissociation-temperature range extending from 60°C to 95°C (0.5°C per 5 s increments).

### Visualization and quantification of EBV positive B-lymphocytes using EBER-ISH

EBER-ISH was performed using the ViewRNA Cell Plus Assay (Affymetrix, USA) according to the manufacturer’s protocol. The EBERs probe set was designed by custom service and ordered from AH Diagnostics. Briefly, fixed suspension cells were spread on poly-L-lysine coated slides in a 96-well plate. The Cell Plus Probe Solution was prepared by diluting Probe Sets 1:100 in pre-warmed Cell Plus Probe Set Diluent and vortexed briefly. The cells were overlaid with Cell Plus Probe Solution (400 μl per well) and gently rocked to mix and distribute the diluted target probe for 2 h at 40°C. Next, we aspirated the Cell Plus Probe Solution and gently and extensively washed the cells with the Cell Plus RNA Wash Buffer Solution using a dropper or pipette to slowly and carefully add 800 μl per well. The cells were covered with Wash Buffer Solution for 24 h at 4°C in the dark. The next day, the samples were pre-warmed to room temperature. The Cell Plus RNA Wash Buffer Solution was aspirated, and the cells were overlaid with Cell Plus Amplifier Diluent (400 μl with 15 μl Cell Plus PreAmplifier Mix) for 1 h at 40°C. The cells were washed extensively, and hybridized with label probes, followed by developing with Fast Red Substrate. Finally, cells were counterstained with DAPI on the slides and mounted with Antifade Reagent (Invitrogen), and imaged at 10X and 20X magnification using Zeiss Axio Imager Z1. Images were taken from six fields and checked for EBERs staining manually.

### Single-cell preparation and sequencing library construction

Two blood samples were obtained in the Department of Clinical Microbiology at Sahlgrenska University Hospital. PBMC were isolated from whole blood using Lymphoprep (Stemcell). B-lymphocytes were enriched using Dynabeads Untouched Human B Cells Kit (Invitrogen). Each sample was loaded into two lanes (i.e. aiming at 20,000 cells per sample). Single-cell gene expression libraries and Single-cell BCR V(D)J libraries were prepared following the 10x user guide CG000186 Rev A (5’ version 1) including doublet removal according to 10x instructions. The sequencing in brief, the library was pooled and sequenced on the NovaSeq6000 platform sequenced on the NovaSeq6000 platform using a single SP flow cell with two lanes, aiming for 27,500 and 5000 sequencing reads per cell for the Gene Expression library and the VDJ library, respectively. One of the libraries from the EBV patient was subjected to an additional sequencing round. The data from all sequencing libraries for each patient were merged during the analysis.

### Single-cell data preprocessing

The raw single-cell sequencing datasets were aligned to the human genome Grch38 with an added EBV reference genome NC_007605.1 as an extra chromosome. To increase sensitivity another alignment was performed using Grch38 with a modified EBV genome in which the genes were removed and the entire genome was considered as a single gene. For the alignment and calculation, the 10x cellranger version 3.0.2 was used. The total cell number and sequencing saturation was analyzed by Cellranger using the merged datasets for each sample. A filtered matrix was generated from the Cellranger count function using standard parameters. EBV statistics were based on the filtered matrix, and the viral transcripts were annotated manually based on the mappling location and strand information in the bam file. The fused EBV genome is available in Github (https://github.com/TangLabGOT/Reference-Genomes).

### Clonality analysis

The raw single-cell VDJ datasets were aligned to the GRCh38 VDJ genes using the `cellranger VDJ` function from cellranger version 3.0.2 with standard parameters. Using the clonotype info summary file the clonal expansion factor was calculated through Equation 1, where *x_i_* =Total number of cells in clonotype *i*, *n* =Total number of clonotypes, and *m* = Total number of cells.

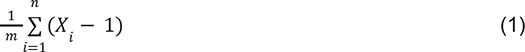

Once each of the cells had been designated a VDJ-profile, each cell was also categorized into a clonotype. After determining the actual EBV positive cells based on the gene expression analysis, the VDJ clonotype analysis was used to infer a positive EBV status of each cell that shared the same CDR3 sequence, i.e. was determined to be the same clonotype, as an actual EBV positive cell. This combination of gene expression analysis and VDJ analysis was key in determining the inferred EBV positive cells as was performed using an in-house python script.

### Single-cell analysis

Single-cell analysis can be divided into four separate parts:

A. No integration, EBV patient and control single-cell analysis
B. Integration, public control single-cell analysis
C. Integration, EBV patient and control single-cell analysis
D. Integration, EBV patient, control, public control, DLBCL, and LCL single-cell analysis

#### A. No integration, EBV patient and control single-cell analysis

Filtered matrix from the Cellranger output was processed using the R package Seurat (4.1.0). Droplets with less than 200 genes were filtered out. Cells with less than 200 features, more than 9000 features, or mitochondrial RNA percentage more than 20% were excluded from further analysis. Normalization was performed using the `NormalizeData` function with no additional parameters. Variable features were found using the `FindVariableFeatures` function with selection methods “vst” and a threshold of 3500 features as parameters. Rescaling and dimensionality reduction was achieved using the `ScaleData` and `RunPCA` function, respectively, using standard settings. Clustering was done through the use of the `FindNeighbors` function using the first ten dimensions in conjunction with the `FindClusters` function with a resolution of 0.5. Data visualization was done using the `RunUMAP` function that in turn uses the UMAP data visualization algorithm while using the ten first dimensions. Based on the visualization and the expression of CD27, IgD, and IgM, each cluster were relabeled into one of four categories: IgD/IgM-Memory B-lymphocytes, IgD/IgM+ Memory B-lymphocytes, Naïve B-lymphocytes, Unassigned. Using the VDJ and clonality data, each cellular barcode that was flagged by the VDJ analysis to be of the same clonal expansion as an EBV infected clonotype was relabeled in the UMAP visualization as “EBV+”. Differential Gene Expression analysis was performed using the `FindAllMarkers` function with parameters: only.pos = True, min.pct = 0.25, logfc.threshold = 0.25.

#### B. Integration, public control single-cell analysis

The publically available healthy B-lymphocyte single-cell data was first merged and integrated according to Seurat’s instruction when performing “Fast integration using reciprocal PCA (RPCA)”. Once integrated, scaling was done using the ScaleData function with the 2000 variable genes found using the FindVariableFeatures function. Once scaled, PCA was performed using the top 20 most variable PCs while also specifying the variable feature to be the above-mentioned 2000 genes, however, with all but the IGHD and IGHM immunoglobulin genes filtered. UMAP visualization was performed using the `RunUMAP` function including the first 20 dimensions. Clustering was performed using the `FindNeighbors` and `FindClusters` function with special parameters first 20 dimensions and resolution of 0.9 for respective functions. Based on the visualization and the expression of CD27, IgD, and IgM, each cluster were relabeled into one of four categories: IgD/IgM-Memory B-lymphocytes, IgD/IgM+ Memory B-lymphocytes, Naïve B-lymphocytes, Unassigned. The IgD/IgM-memory B-lymphocyte population was extracted and Differential Gene Expression was performed comparing each respective sample (N21, N22, and N23) with the other two samples (N22+N23, N21+N23, N21+N22) using the FindAllMarkers function with parameters: only.pos = False, min.pct = 0, logfc.threshold = 0. A heuristic filter was implemented (|Average Log2FC| ≥ Log2(1.1), Adjusted P-value ≤ 0.05) to filter out genes with either too small of an effect or that were not significant enough. Once this curated list had been generated, this list was used to filter out overlapping, individual-based differences in expression pattern for (C.) analysis.

#### C. Integration, EBV patient and control single-cell analysis

The EBV patient single-cell data and the control single-cell data was merged, integrated and processed in a similar manner as the (B.) analysis up until after the `FindNeighbors` and `FindClusters` function. Based on the visualization and the expression of CD27, IgD, and IgM, each cluster were relabeled into one of four categories: IgD/IgM-Memory B-lymphocytes, IgD/IgM+ Memory B-lymphocytes, Naïve B-lymphocytes, Unassigned. Using the clonality analysis, each cellular barcode that was flagged by the VDJ analysis to be of the same clonal expansion as an EBV infected cell was relabeled in the UMAP visualization as “EBV+”. Differential Gene Expression was performed twice comparing the inferred EBV+ lymphocytes against the control’s IgD/IgM-memory B-lymphocytes as well as comparing the EBV patient’s naïve B-lymphocytes against the control’s naïve B-lymphocytes. This comparison was performed using the FindAllMarkers function with parameters: only.pos = False, min.pct = 0, logfc.threshold = 0. A heuristic filter was implemented (|Average Log2FC| ≥ Log2(1.1), Adjusted P-value ≤ 0.05) to filter out genes with either too small of an effect or that were not significant enough. Once these two curated lists had been generated, the list of differentially expressed genes from the naïve B-lymphocytes were used as a filter to remove genes based on the innate difference in expression pattern between the EBV patient and the control. Once filtered, the curated list of differentially expressed genes from (B.) analysis was used to further filter individual based differences, as aforementioned. This list of double curated genes were then split into two separate lists depending on if the gene had an increased or decreased level of expression. These two lists were then analyzed for biological processes pathway enrichment using GeneOntology Enrichment Analysis with parameters: Fisher’s exact test, FDR as multiple testing correction, and a significance threshold of adjusted p-value ≥ 0.05.

#### D. Integration, EBV patient, control, public control, DLBCL, and LCL single-cell analysis

The EBV patient, control, N21-N23 public control, C01-C03 public LCL, and four DLBCL single-cell data were merged, integrated, and processed in a similar manner as the (B.) analysis, up until the `RunPCA` function. The `RunPCA`, the `FindNeighbors`, and the `RunUMAP` function was run with the first 30 dimensions as the parameters while the `FindClusters` was run with 0.9 as resolution. Each cluster’s original dataset composition was displayed in percentage. The clusters with cells equal to or above 50 percent were classified as DLBCL/LCL. Each cluster’s differentially expressed genes were generated through the FindAllMarkers function with parameters: only.pos = False, min.pct = 0, logfc.threshold = 0. Using this list of differentially expressed genes, cluster 15 and cluster 17 were deemed as being tumor infiltrating lymphocytes and were thus excluded from further analysis, leaving cluster 4, 8, 11, 13, and 14 to be relabeled as “DLBCL/LCL”. As previously described, each cellular barcode that was flagged by the VDJ analysis to be of the same clonal expansion as an EBV infected cell was relabeled in the UMAP visualization as “EBV+”.

### Sequencing context and metadata analysis

EBV patient, the control, C01-03, and N21-23 were aligned to the fused annotated EBV reference genome^23^. EBV genes per cell was calculated based on the alignment results. Due to limitation in accessing the raw DLBCL sequencing data, the number of EBV genes per cell was set to zero as DLBCL is rarely associated with EBV. Using the seurat package as described previously, the mRNA content of every cell was extracted and ordered in an ascending manner based on mRNA per cell and total number of EBV genes expressed in the cell.

## Acknowledgements

This project was supported by funding from Region Västra Götaland, the Assar Gabrielssons Research Foundation and the Swedish Society for Medical Research (S21-0083). We thank the Bioinformatics Core Facility at the Sahlgrenska Academy for bioinformatics analyses. The computations and data handling were enabled by resources provided by the Swedish National Infrastructure for Computing (SNIC, project sens2018120) at Uppsala Multidisciplinary Center for Advanced Computational Science (UPPMAX) partially funded by the Swedish Research Council through grant agreement no. 2018-05973. The sequencing was conducted by NGI Sweden. The study design and methods including waiver of informed consent were approved by the Swedish Ethical Review Authority, Region Göteborg (191-18).Included samples from patients with from the Department of Clinical Microbiology at Sahlgrenska University Hospital (Biobank VDB 263).The study design and methods including waiver of informed consent were approved by the Swedish Ethical Review Authority, Region Göteborg (191-18).

## References

1. Moustafa A, et al. The blood DNA virome in 8,000 humans. PLoS Pathog 13, e1006292 (2017).

2. Hsu JL, Glaser SL. Epstein-barr virus-associated malignancies: epidemiologic patterns and etiologic implications. Crit Rev Oncol Hematol 34, 27–53 (2000).

3. Lyu M, et al. Dissecting the Landscape of Activated CMV-Stimulated CD4+ T Cells in Humans by Linking Single-Cell RNA-Seq With T-Cell Receptor Sequencing. Front Immunol. 12, (2021).

4. Chakravorty S, et al. Integrated Pan-Cancer Map of EBV-Associated Neoplasms Reveals Functional Host-Virus Interactions. Cancer Res 79, 6010–6023 (2019).

5. Farrell PJ. Epstein-Barr Virus and Cancer. Annu Rev Pathol 14, 29–53 (2019).

6. Liu Y, et al. Tumour heterogeneity and intercellular networks of nasopharyngeal carcinoma at single cell resolution. Nat Commun 12, 741 (2021).

7. Gong L, et al. Comprehensive single-cell sequencing reveals the stromal dynamics and tumor-specific characteristics in the microenvironment of nasopharyngeal carcinoma. Nat Commun 12, 1540 (2021).

8. Jin S, et al. Single-cell transcriptomic analysis defines the interplay between tumor cells, viral infection, and the microenvironment in nasopharyngeal carcinoma. Cell Res 30, 950–965 (2020).

9. Chen YP, et al. Single-cell transcriptomics reveals regulators underlying immune cell diversity and immune subtypes associated with prognosis in nasopharyngeal carcinoma. Cell Res 30, 1024–1042 (2020).

10. SoRelle ED, et al. Single-cell RNA-seq reveals transcriptomic heterogeneity mediated by host-pathogen dynamics in lymphoblastoid cell lines. Elife 10, (2021).

11. Lanz TV, et al. Clonally expanded B cells in multiple sclerosis bind EBV EBNA1 and GlialCAM. Nature 603, 321–327 (2022).

12. Kurth J, Hansmann ML, Rajewsky K, Kuppers R. Epstein-Barr virus-infected B cells expanding in germinal centers of infectious mononucleosis patients do not participate in the germinal center reaction. Proc Natl Acad Sci U S A 100, 4730–4735 (2003).

13. Tang KW, Alaei-Mahabadi B, Samuelsson T, Lindh M, Larsson E. The landscape of viral expression and host gene fusion and adaptation in human cancer. Nat Commun 4, 2513 (2013).

14. Zapatka M, et al. The landscape of viral associations in human cancers. Nat Genet 52, 320–330 (2020).

15. Tang KW, Hellstrand K, Larsson E. Absence of cytomegalovirus in high-coverage DNA sequencing of human glioblastoma multiforme. Int J Cancer 136, 977–981 (2015).

16. Tang KW, Larsson E. Tumour virology in the era of high-throughput genomics. Philos Trans R Soc Lond B Biol Sci 372, (2017).

17. Chan KC, et al. Molecular characterization of circulating EBV DNA in the plasma of nasopharyngeal carcinoma and lymphoma patients. Cancer Res 63, 2028–2032 (2003).

18. Bjornevik K, et al. Longitudinal analysis reveals high prevalence of Epstein-Barr virus associated with multiple sclerosis. Science 375, 296–301 (2022).

19. Kanakry JA, et al. The clinical significance of EBV DNA in the plasma and peripheral blood mononuclear cells of patients with or without EBV diseases. Blood 127, 2007–2017 (2016).

20. Strong MJ, et al. Differences in gastric carcinoma microenvironment stratify according to EBV infection intensity: implications for possible immune adjuvant therapy. PLoS Pathog 9, e1003341 (2013).

21. Green M, Michaels MG. Epstein-Barr virus infection and posttransplant lymphoproliferative disorder. Am J Transplant 13 Suppl 3, 41–54; quiz 54 (2013).

22. Andersson D, et al. Properties of targeted preamplification in DNA and cDNA quantification. Expert Rev Mol Diagn 15, 1085–1100 (2015).

23. Ziegler P, et al. A primary nasopharyngeal three-dimensional air-liquid interface cell culture model of the pseudostratified epithelium reveals differential donor- and cell type-specific susceptibility to Epstein-Barr virus infection. PLoS Pathog 17, e1009041 (2021).

24. Holmqvist I, Backerholm A, Tian Y, Xie G, Thorell K, Tang KW. FLAME: long-read bioinformatics tool for comprehensive spliceome characterization. RNA 27, 1127–1139 (2021).

25. von Budingen HC, et al. B cell exchange across the blood-brain barrier in multiple sclerosis. J Clin Invest 122, 4533–4543 (2012).

26. Grande BM, et al. Genome-wide discovery of somatic coding and noncoding mutations in pediatric endemic and sporadic Burkitt lymphoma. Blood 133, 1313–1324 (2019).

27. Souza TA, Stollar BD, Sullivan JL, Luzuriaga K, Thorley-Lawson DA. Influence of EBV on the peripheral blood memory B cell compartment. J Immunol 179, 3153–3160 (2007).

28. Mrozek-Gorska P, et al. Epstein-Barr virus reprograms human B lymphocytes immediately in the prelatent phase of infection. Proc Natl Acad Sci U S A 116, 16046–16055 (2019).

29. Niesters HG, van Esser J, Fries E, Wolthers KC, Cornelissen J, Osterhaus AD. Development of a real-time quantitative assay for detection of Epstein-Barr virus. J Clin Microbiol 38, 712–715 (2000).

30. Kullberg-Lindh C, Olofsson S, Brune M, Lindh M. Comparison of serum and whole blood levels of cytomegalovirus and Epstein-Barr virus DNA. Transpl Infect Dis 10, 308–315 (2008).

