## Supplemental Table 4 for "Detection of latent Epstein-Barr virus gene expression in single-cell sequencing of peripheral blood mononuclear cells"

| **Clone** | **Heavy chain** | | | | **Somatic hypermutations in VH** | | | | | | **Light chain** | | |
|---|---|---|---|---|---|---|---|---|---|---|---|---|---|
| | V | D | J | C | CDR1 | CDR2 | CDR3 | FR1 | FR2 | FR3 | κ/λ | V | J |
| 1 | 1-69 | 2OR15-2B | 3 | A1 | 9 | 4 | 16 | 2 | 0 | 5 | κ | 3-20 | 1 |
| 2 | 4-4 | 4-17 | 4 | A1 | 5 | 4 | 10 | 1 | 0 | 3 | κ | 3-20 | 4 |
| 3 | 3-7 | 3-10 | 6 | G1 | 4 | 0 | 13 | 1 | 0 | 2 | λ | 1-40 | 2 |
| 4 | 1-69D | 3-16 | 1 | A1 | 6 | 4 | 13 | 4 | 2 | 3 | κ | 3-11 | 2 |
| 5 | 1-69D | 3OR15-3A | 6 | G1 | 4 | 6 | 7 | 1 | 1 | 4 | κ | 2-28 | 2 |
| 6 | 2-70 | 3-16 | 6 | G1 | 8 | 1 | 14 | 0 | 0 | 0 | λ | 3-21 | 2 |
| 7 | 1-69D | 5-18 | 6 | A1 | 3 | 4 | 10 | 3 | 1 | 4 | λ | 2-23 | 2 |
| 8 | 1-3 | 7-27 | 3 | G3 | 6 | 6 | 9 | 3 | 3 | 5 | κ | 3-11 | 4 |
| 9 | 1-2 | 3-9 | 1 | G1 | 1 | 3 | 12 | 1 | 2 | 2 | κ | 3-20 | 4 |
| 10 | 4-39 | 5-12 | 4 | G1 | 6 | 5 | 6 | 1 | 2 | 5 | λ | 2-8 | 2 |
